# Innovative approach for vaccinating Nile tilapia, *Oreochromis niloticus* against *Streptococcus agalactiae* using an ozone nanobubble pre-treatment, VAC in BAG and VAC in FEED

**DOI:** 10.1101/2023.06.09.544435

**Authors:** Nguyen Tien Vinh, Ha Thanh Dong, Saengchan Senapin, Suntree Pumpuang, Nguyen Giang Thu Lan, Bulakorn Wilairat, Pradnya R. Garud, Sophie St-Hilaire, Nguyen Vu Linh, Wattana Phanphut, Andrew P. Shinn

**Author notes:** Corresponding author: H. T. Dong.

## Abstract

The treatment of Nile tilapia with ozone nanobubbles (ONb) prior to vaccination with an immersible heat-killed *Streptococcus agalactiae* (*Sa*) vaccine has been reported to modulate and enhance both innate and specific immunity. The efficacy of this novel vaccination strategy is explored further in field trials. This strategy involved a short-term treatment of ONb to activate the fish’s immunity prior to immersion vaccination during their transportation in oxygenated plastic bags (VAC in BAG), followed by two oral boosters during the grow-out stage mixing vaccine in feed (VAC in FEED). The field trial was conducted over 112 days in open cages, comprising four groups: normal aeration control (AC), normal aeration + vaccine (AV), ONb control (NC), and ONb + vaccine (NV). The efficacy of the vaccine was evaluated by measuring specific antibodies for *S. agalactiae*, monitoring expressions of *IgM* and *IgT* transcripts in the gills and head kidney every two weeks, and a laboratory pathogen challenge. Results found that fish in the NV group had significant increases in anti-*S. agalactiae* antibodies after the primary dose, whereas fish in the AV group required an oral booster dose to produce significant anti-*S. agalactiae* antibodies. In the vaccinated groups (AV and NV), only *IgM* was observed to be upregulated at 14 days post-immersion (dpi), while this gene was upregulated in both gills and head kidney in the NC group. No statistically significant upregulation of *IgT* was recorded in any group at any time point. Despite a decline in the levels of specific antibodies among the vaccinated groups at the time of challenge (88 dpi), the NV and AV groups demonstrated a relative percent survival (RPS) of 50% and 46.7%, respectively, following a relatively high injection dose of *S. agalactiae* injection (0.1 mL of 10^8^ CFU/mL). In summary, this ONb, VAC in BAG and VAC in FEED vaccination strategy represents a promising alternative to the undesirable handling and costly injection approach used within the Nile tilapia industry.

## 1. Introduction

Nile tilapia (*Oreochromis niloticus*) holds significant commercial value as one of the top three cultured freshwater fish species (FAO, 2020). Various farming systems are employed for tilapia, including pond, cage, tank, raceway, and aquaponic systems (Romana-Eguia et al., 2020). The rapid growth of an intensive cage culture system is attributable to its simple operational procedures and favorable economic viability (Chitmanat et al., 2016). Episodes of disease, however, frequently follow events of high stocking density, poor water conditions (such as marked changes in temperature, low dissolved oxygen, or elevated ammonia concentrations, etc.), climate change, and improper handling (Georgiadis et al., 2001; Amal et al., 2015). Disease outbreaks and subsequent economic loss are a major impediment to the sustainable growth of the tilapia industry. Among these, streptococcal infections are commonly reported; outbreaks of *Streptococcus agalactiae* in cage culture can result in significant loss (Mian et al., 2009; Jantrakajorn et al., 2014; Shinn et al., 2023).

Utilizing vaccination as a means of addressing infectious diseases in aquaculture and in mitigating the proliferation of antimicrobial resistance (AMR) holds promising potential, as perceived in terrestrial farmed animals and humans (Henriksson et al., 2017; Jansen & Anderson, 2018; Micoli et al., 2021). To date, most commercially available vaccines for fish are either in an immersible or injectable form. The latter is favored by the salmon industry as they tend to provide good immunity (Ma et al., 2019; Miccoli et al., 2021). However, injection vaccines are expensive to administer so they are less appealing for fish with low market value. In addition, there are significant challenges associated with the injection method, notably in the labor-intensive nature and the logistics of vaccinating fish that have already been stocked in grow-out systems. For those reasons, alternative vaccine delivery modes, such as immersion and oral, are sought as more practically applicable in fish farms. Immersion and oral vaccines against *S. agalactiae* have been the topic of several recently published articles (Zhu et al., 2017; Abu-Elala et al., 2019; Yao et al., 2019; Ke et al., 2021;), however, mucosal application is still being conceived and limited to lab scale experiments.

Although there are benefits to the application of both immersion and oral vaccines in the field, i.e., logistically they are easier to administer, they negate the need for handling and consequentially result in lower levels of stress in the fish population, however, they may elicit inferior levels of immunity when compared to those delivered by injection (Brudeseth et al., 2013; Bøgwald & Dalmo, 2019; Mondal & Thomas, 2022). Ensuring the efficacy of a mucosal vaccine, therefore, may require their application with adjuvants or with treatments to obtain an appropriate immune response from the fish (Adams, 2019; Bøgwald & Dalmo, 2019). Recent research has found that the use of an ozone nanobubble (ONb; typically less than 130 nm) treatment can modulate the innate immune response of fish (Linh et al., 2021) and enhance the efficacy of an immersible *S. agalactiae* vaccine, making it a promising application for use in aquaculture (Linh et al., 2022). To prolong the vaccine efficacy, a booster dose is recommended during the grow-out culture period, and oral vaccines are considered more appealing over other delivery methods (Plumb & Vinitnantharat, 1993; Chettri et al., 2015; Schmidt et al., 2016).

To address and mitigate issues associated with infectious diseases in the Nile tilapia aquaculture industry, it is necessary to promote a locally enabled, effective and convenient vaccine application program. In this research, we investigated a new vaccination strategy to prevent *S. agalactiae* infection in cage cultured tilapia, with a focus on the ease of conducting vaccination by immersion, and oral administration. Here we refer to the technique as “an ozone nanobubble pre-treatment”, “VAC in BAG” and “VAC in FEED”. This approach encompasses pre-treatment with ONb to bolster the fish’s immune response before immersion vaccination during their transportation in a bag (referred to as VAC in BAG) and the subsequent administration of oral booster doses (known as VAC in FEED) during the grow-out phase. With this novel procedure, we hope to create a game changing solution to conventional vaccination ushering in a novel strategy for the use of vaccines in aquaculture.

## 2. Materials and methods

### 2.1 Experimental design

A total of 3200 healthy, juvenile Nile tilapia, *O. niloticus* (4.77 ± 0.96 g), were obtained from the Asian Institute of Technology, Thailand, and used for the vaccine field trial. Fish were segregated into four groups: (1) normal aeration - control (AC) which was not treated with either ozone nanobubbles nor vaccine, (2) ozone nanobubbles - control (NC) where the fish were treated with ozone nanobubbles but not exposed to the vaccine, (3) normal aeration + vaccine (AV) where the fish were immunized with the vaccine after normal aeration, and (4) ozone nanobubbles + vaccine (NV) where the fish were treated with ONb prior to their immersion in the vaccine.

### 2.2 Husbandry practice

The grow out site selected for the field trial was a ca. 1 hectare pond (∼4 m depth) connected to a nearby canal. The fish were stocked into a cage system comprised of four compartments; each measuring 4 × 2 × 1.5 m (Figure S1). Initial stocking density was 0.3 kg/m^3^ with 800 fish stocked into each compartment. A line of solar panel powered paddle wheels were positioned close to the edge of the two cage compartments stocked with the AC and NC test groups of fish to provide oxygen for fish. In addition to the experimental fish, the pond was inhabited by wild population of tilapia that had established from escapees from previous farm production cycles. A plastic net was employed to provide shading, thereby mitigating excessive sunlight and heat while simultaneously serving as a deterrent against avian intrusions. Fish were fed twice daily (Lee Feed Mill Public Company Limited - Win 32, 32% crude protein) at 3% biomass.

### 2.3 Vaccine preparation

The *Sa* vaccine was prepared using *S. agalactiae* strain 2809, isolated from an earlier infection in Nile tilapia (Jhunkeaw et al., 2021). The isolate was initiated in 20 mL of tryptic soy broth (TSB, Becton, Dickinson and Company, USA) at 30°C for 18 hours and then transferred and grown on for another 18 hours in 500 mL TSB broth. After reaching an OD_600_ nm of 1.0, the culture exhibited a concentration of ∼10^9^ CFU/mL.

The vaccine was prepared using a conventional heating method. Following culture, the whole bacterin was subjected to a heat treatment of 60°C for 60 minutes to achieve inactivation. Inactivation was verified by spreading a sample of the vaccine onto agar plates (Tryptic soy agar - TSA). The vaccine was then stored at 4°C (within three days) until required for immersion vaccination.

For the oral booster, the diet was prepared by mixing 100g of the commercial tilapia feed with 20 mL of the *Sa* vaccine and then coating the feed with 10 mL palm oil (Patum Vegetable Oil Co., Ltd., Thailand). The final concentration of the vaccine in the feed was ∼ 2 × 10^8^ CFU/g feed. The feed was then dried at 60°C in an oven for 12 hours, followed by cooling and storing at 4°C. To uphold the quality of the feed-based vaccine presented to fish, the VAC in FEED was prepared 2 days prior to its use in the field. The diet for fish groups without vaccine (AC and NC) during oral booster periods was prepared in the same manner, by substituting the vaccine with TSB of the same amount.

### 2.4 Vaccination procedure

The field vaccination trial is summarized in Figure 1. The fish were starved for 24 hours before vaccination. The first step of the procedure was to treat the tilapia by exposure to either ONb (NC and NV groups) or to normal aeration (AC and AV) for 15 minutes. In each group, the fish were divided into two replicate tanks, with each 200 L tank containing 400 fish in 180 L of freshwater. In both the NC and NV groups, ONb were created using a nanobubble generator (aQua+, Singapore) that was supplied with ozone gas (ozone generator, model CCba15, Coco Technology Co., Ltd.) converted from ∼95% saturated oxygen (oxygen concentrator, model VH5-N, Shenyan Canta Medical Tech. Co., Ltd.) (see the system setup provided in Jhunkeaw et al., 2021). The fish were stocked directly in the tank with the ozone nanobubble generator running. The oxidation-reduction potential (ORP) measured from a multiprobe (YSI Professional Plus, YSI Incorporated, USA) was used to indirectly assess the creation of the ONb. The ORP readings peaked at approximately 700 - 800 mV after 15 minutes, indicating the optimal creation of ONb. The fish were then allowed to acclimate and to stabilize under normal aeration conditions for two hours before being packed for transportation in 4.5 L capacity bags. The water enriched with oxygen nanobubbles was used for the NC and NV groups, while normal water without oxygen nanobubbles was used for AC and AV groups. All bags were filled with oxygen gas before packing for transportation.

**Figure 1.**
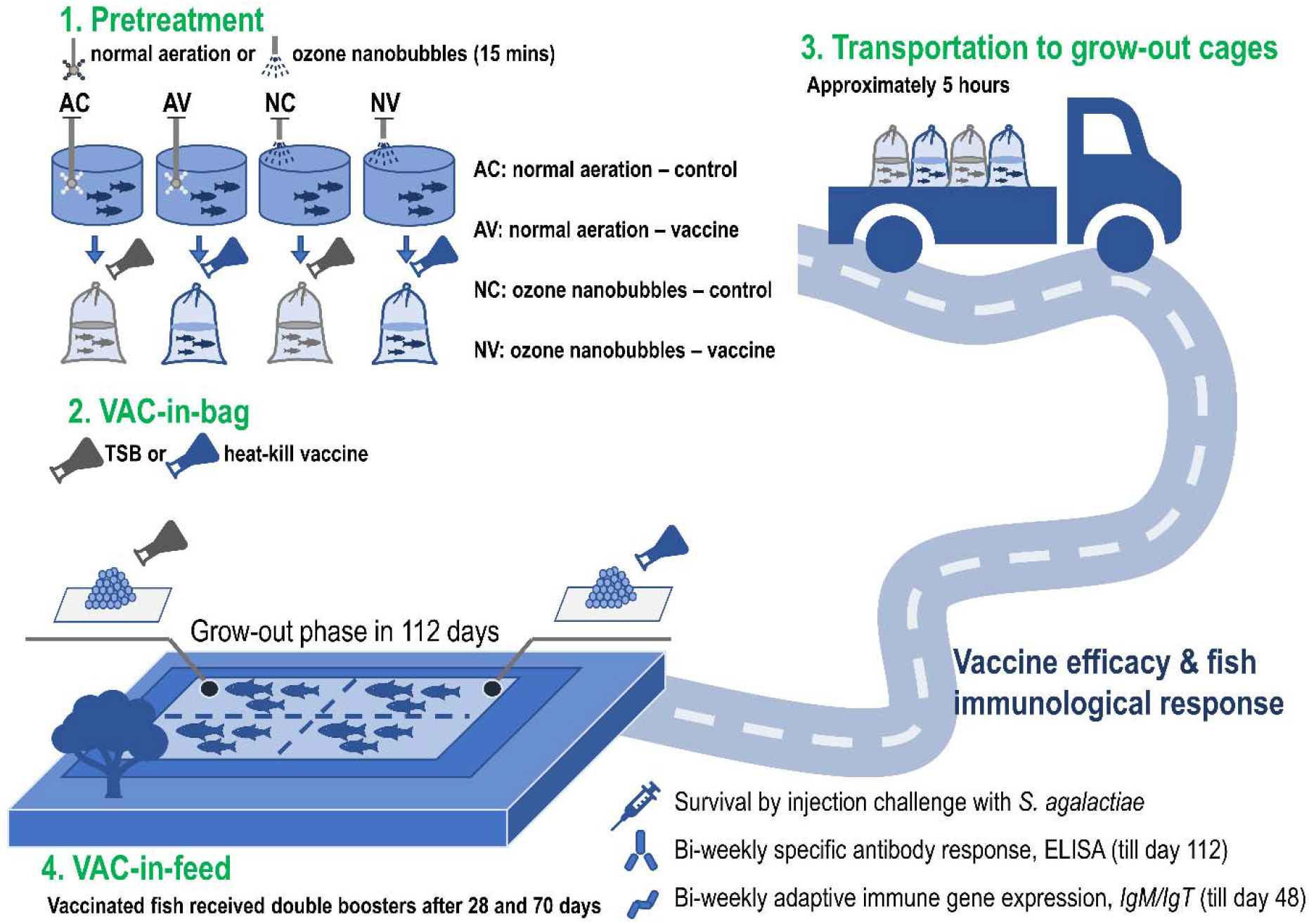
Experimental design of ozone nanobubble pre-treatment, VAC in BAG and VAC in FEED strategy against *Streptococcus agalactiae* in a Nile tilapia, *Oreochromis niloticus*, farm.

Immersion immunization was performed in plastic bags with a capacity of 4.5 L, accommodating 100 fish per bag (i.e., ca. 106 g/L fish biomass), in accordance with the usual stocking densities used for the transportation of juvenile tilapia. In the vaccinated groups (i.e., AV and NV), the heat-killed *S. agalactiae* vaccine (10^9^ CFU/mL) was added at 1% (v/v), while in the non-vaccinated groups (i.e., AC and NC), the vaccine was substituted with an equivalent volume of TBS solution (i.e., 450 mL/bag). After packing, the fish were promptly transported to the grow-out site, a journey that took approximately 5 hours.

During the 112-day grow-out period in the cage-in-pond system, the fish in vaccinated groups (AV and NV) were given two oral vaccine boosters (VAC in FEED) on two periods, i.e., 28-32 and 70-72-days post-immersion (dpi). On each date, the fish were fed at 3% body weight (based on a random sampling of ten fish per treatment group prior to presenting the fish with the VAC in FEED). For the first booster on day 28, the fish were fed the VAC in FEED for five days, while for the second booster given on day 70 the fish were given the VAC in FEED for three consecutive days. Feed mixed with TSB was applied to the control groups (i.e., AC and NC) during the VAC in FEED phases.

### 2.5 Sample collection

Fish sampling was conducted every two weeks starting from day 14 post-immersion. At each time point, 10 fish from each group were randomly selected and processed. Sera samples for an enzyme linked immunosorbent assay (ELISA) were collected until the end of the trial on day 112. Samples of gill and head kidney tissue, however, were collected only until day 42. From day 56 dpi, a non-lethal blood sample was taken from the fish while tissues for RNA were collected from others at post-mortem. For the non-lethal blood sampling, the tilapia were anesthetized in a clove oil (0.5 mL/L) bath for 30 seconds. A blood sample of 200 – 300 µL was collected from the caudal peduncle, and then centrifuged at 6,000 g for 15 minutes to separate the sera; this was then collected and stored at -20 °C for ELISA. For the fish that were processed post-mortem, following termination, a blood sample was taken and processed as described above. Then the first two gill arches from one side were removed followed by a sample of the head kidney for immune gene expression analysis. The tissues were fixed and stored in RNAlater at -20 °C until required for RNA analysis.

### 2.6 Immunoassay of antibody response post-vaccination

The antibody-ELISA assay was conducted following the protocol as described by Linh et al. (2022) using flat-bottom microplates (Costar®, Corning Inc., USA). Briefly, the *Sa* antigen was first coated at ∼10^8^ CFU/mL overnight. The antigen was prepared from the prepared vaccine described above, by collecting bacterial cells from the heat-killed vaccine centrifuged at 2900 g in 15 minutes, following dilution in phosphate buffered saline (1X PBS). Fish sera were diluted in PBSTM (i.e., PBS 1X + 0.05% Tween-20 + 5% skimmed milk) at a ratio of 1:511. Primary (anti-mouse *IgM* tilapia) and secondary (anti-mouse antibody horseradish peroxidase conjugate) antibodies were subsequently added at ratios of 1:199 and 1:2999 respectively, each diluted in PBSTM. A chromogenic substrate, 3,3’, 5,5’-tetramethylbenzidine (TMB) was added to signalize antibody binding. The reaction was stopped using 100 µL of 2M H_2_SO_4_ before the samples were read on a microplate reader (AC3000, Azure Biosystems, USA) at a wavelength of 450 nm (OD_450_).

### 2.7 Gene expression analysis

Total RNA extracted from the gills and head kidneys were used for the specific-immune gene expression analysis. Immunoglobulin transcripts encoding *IgM* and *IgT* were selected as target genes, by using the same primer pairs and quantitative PCR (qPCR) conditions as described previously in Linh et al. (2021, 2022). Relative gene expression was performed by the 2^-ΔΔCt^ method (Livak & Schmittgen, 2001), using the *EF1*α expression of the AC group at the same sampling point for normalization.

### 2.8 Experimental challenge with *Streptococcus agalactiae*

On day 84 of the trial, 120 fish (30 fish per group; av. wt. 80-90 g per fish) that had been pre-starved for 24 h were translocated from the field site to a biosecure disease challenge facility for the experimental challenge with *S. agalactiae*. On arrival at the facility, the fish were allocated to 200 L static, aerated, polyethylene tanks filled with 150 L freshwater and allowed to acclimate for 3 days prior to challenge. During the acclimation period, the feeding regime was gradually reduced to 1% body weight day^-1^. This lower rate of feeding continued up until their experimental challenge with *S. agalactiae*. Following injection, feeding was then withheld. Heavy rates of mortality typically ensue during which the fish do not feed. The water was exchanged at fifty percent every two days during the duration of the challenge. Water quality in the tanks was assessed daily to ensure that parameters remained within acceptable levels for their culture and welfare.

The *Streptococcus* challenge was performed on day 88 of the trial. The pathogen was cultured as described in the ‘Vaccine preparation’ section. The bacteria cells were collected by centrifugation at 2900 g for 15 minutes and then diluted in 1X PBS to 10^8^ CFU/ mL for injection. Each fish was injected with 0.1 mL of the prepared pathogen culture, equivalent to 10^7^ CFU/ fish as described previously (see Linh et al., 2022). The challenge lasted for 15 days, during which, they were monitored at 7.00 am, 12.00 pm midday, 17.00 pm, and 22.00 pm every day, during which the behaviors of the fish including abnormal signs, moribund fish, or mortalities were recorded. Mortalities were immediately removed and any moribund fish were terminated by an overdose of clove oil and then necropsied. The liver and brain tissues of moribund fish were streaked on to TSA plates to recover the pathogen. At the end of the challenge, all survivors were terminated by an overdose of clove oil. All fish carcasses and tissues were autoclaved as part of onsite biosecurity practices.

### 2.9 Statistical analysis

Data exploration and summaries were performed using dplyr ver. 1.0.10 (Wickham et al., 2022). For both the antibody assay and the gene expression analysis, Kruskal-Wallis tests were performed at each sampling point, followed by a Dunn’s Multiple Comparison Test with the Bonferroni *p* adjust method using rstatix ver. 0.7.2 (Kassambara, 2023). The level of significance for all tests was set at a *p*-value < 0.05. Graphs were plotted using the ggplot2 package (Wickham, 2016). For the immunoassay, a cutoff threshold (Frey et al., 1998) was calculated at the upper 95% interval using the OD readings of the AC group at 14 dpi to determine positive- and negative-anti-*Sa* antibody production of individual fish. The challenge test result was expressed as relative percent survival (RPS) according to the mortality of the AC group (1 – [cumulative percent mortality in target group / cumulative percent mortality in AC group] x 100). Cumulative percent survival was presented by a Kaplan-Meier plot, adhering to the analysis of R packages - survival ver. 3.5-0 (Therneau, 2023) and surminer ver. 0.4.9 (Kassambara et al., 2021).

### 2.10 Ethics statement

The experimental procedures conducted in this study were evaluated and approved by the Thai Institutional Animal Care and Use Committee (approval no. MUSC62-039-503). Additionally, the trials were overseen by an aquatic veterinarian.

## 3. Results

### 3.1 Observation of fish health and environmental parameters during the experiment

During the immersion VAC in BAG phase of the study, no abnormalities or mortalities were observed. The tilapia resumed normal schooling and feeding behaviors after stocking into the grow-out ponds. Fish readily accepted the feed when presented with the vaccine dressed feed. Environmental parameters (measured between 11:00 am – 12.00 pm midday) during the field component of the vaccine trial were determined to be 5.72 ± 0.47 mg/L dissolved oxygen (DO); 8.40 ± 0.35 pH units; and, 31.08 ± 0.30 °C temperature.

### 3.2 Specific antibody (IgM) response after vaccination

The average *Sa*-specific antibody production (*IgM*) of the main experimental target group, i.e., the fish vaccinated using an ONb pre-treatment (NV), was consistently higher than the other experimental groups from day 14 to 84 post-immunization (Figure 2). Following immersion in the *Sa* vaccine, only the NV group displayed a significant increase in OD readings compared to the normal aerated-unvaccinated group (AC) at day 14 (OD_450 nm_ = 0.341 ± 0.036) and day 28 (OD_450 nm_ = 0.338 ± 0.026). The OD readings of the vaccinated group without the ONb pre-treatment (AV) was not statistically different from the controls until 28 dpi (OD_450 nm_ = 0.272 ± 0.029).

**Figure 2.**
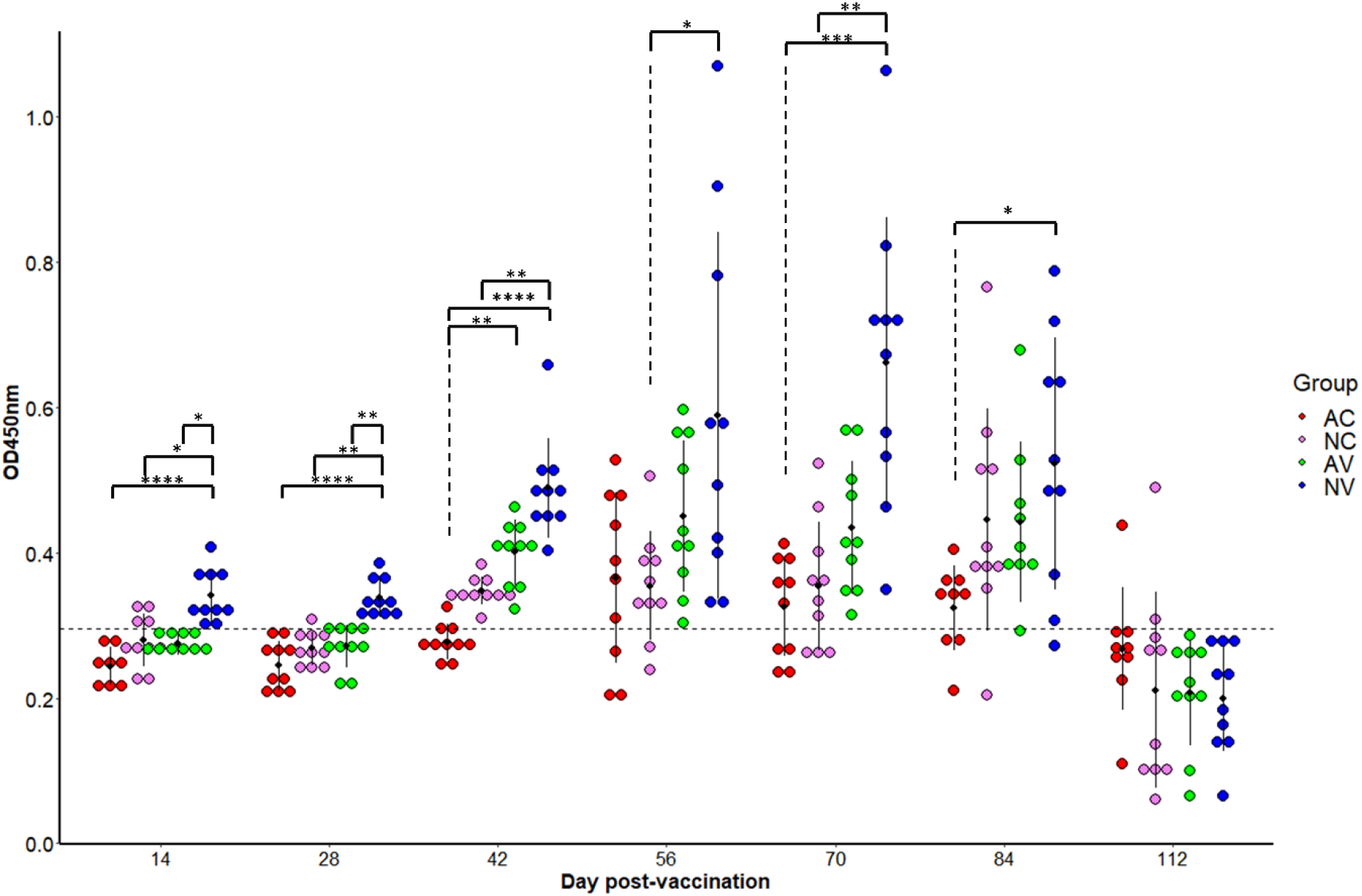
Antibody response of Nile tilapia, *Oreochromis niloticus*, during the 112-day field trial. Each dot represents one individual biological replicate. Error bars indicate average OD_450_ nm with standard deviation. Within the same sampling point, the connector with asterisks indicates a significant difference between two groups using Dunn’s test (*p* adjust with Bonferroni method). Different levels are (*) *p < 0*.*05*, (**) *p < 0*.*01*, (***) *p < 0*.*001*, (****) *p < 0*.*0001*. Cut-off level (dashed line) was calculated based on the significant level at *p* = 0.05, based on the OD readings of *O. niloticus* of the AC group. Abbreviations: AC = normal aeration control; AV = normal aeration + vaccine; NC = ozone nanobubble treatment control; NV = ozone nanobubble treatment + vaccine.

The first vaccine oral booster was administered on days 28-32 (5 days) after the initial immersion. The first significant difference in antibody (Ab) production displayed by the AV group, when compared to the AC group, was on 42 dpi (i.e., OD_450 nm_ = 0.401 ± 0.045); thereafter there was no statistically significant differences in Ab levels between the AV, AC and NC groups. The levels of Ab in the NV group, however, were markedly higher reaching an OD_450 nm_ of 0.489 ± 0.068 at 42 dpi and then peaking significantly at 56 (OD_450 nm_ = 0.589 ± 0.252) and 70 dpi (0.662 ± 0.200). Lower antibody levels in the NV group, however, were measured on day 84, i.e., two weeks following the second oral vaccine booster administered on days 70. Antibody levels, however, remained significantly higher than those determined from fish in the AC group. No significant difference in specific antibody production was observed between the vaccinated groups at 112 dpi.

Based on the cutoff at the upper 95% interval of antibody levels measured in the AC group, most fish in the NV group had positive antibody titers specific to *Sa* throughout the vaccine trial until 84 dpi. Positive antibodies were recorded from day 42 in all sampled fish from the AV group. In the non-vaccinated group (NC), fish began to show positive *Sa*-specific antibodies (i.e., in 100% in those fish that were sampled) on the same day, albeit without a significant difference (*p* > 0.05) when compared to the AC group. Antibodies in many fish from the AC and NC groups exceeded the threshold from day 56. At the end of the period over which antibodies were measured, i.e., trial termination on 112 dpi), most fish displayed antibody levels that were below the established threshold.

### 3.3 Expression of immunoglobulin genes after vaccination

*IgM* expression in the gills of three groups, NC, AV, and NV, was significantly upregulated 14 days after immersion, with average fold changes ranging from above 3.1 to 4.6 (Figure 3). In the head kidney, the highest expression was observed in the NC group, with an approximately six-fold increase. Meanwhile, no significant *IgM* transcripts were obtained in either the AV or the NV vaccinated groups at the same sampling point. At subsequent sampling points, 28 and 42 dpi, there was no significant upregulation in *IgM* expression in any of the groups.

**Figure 3.**
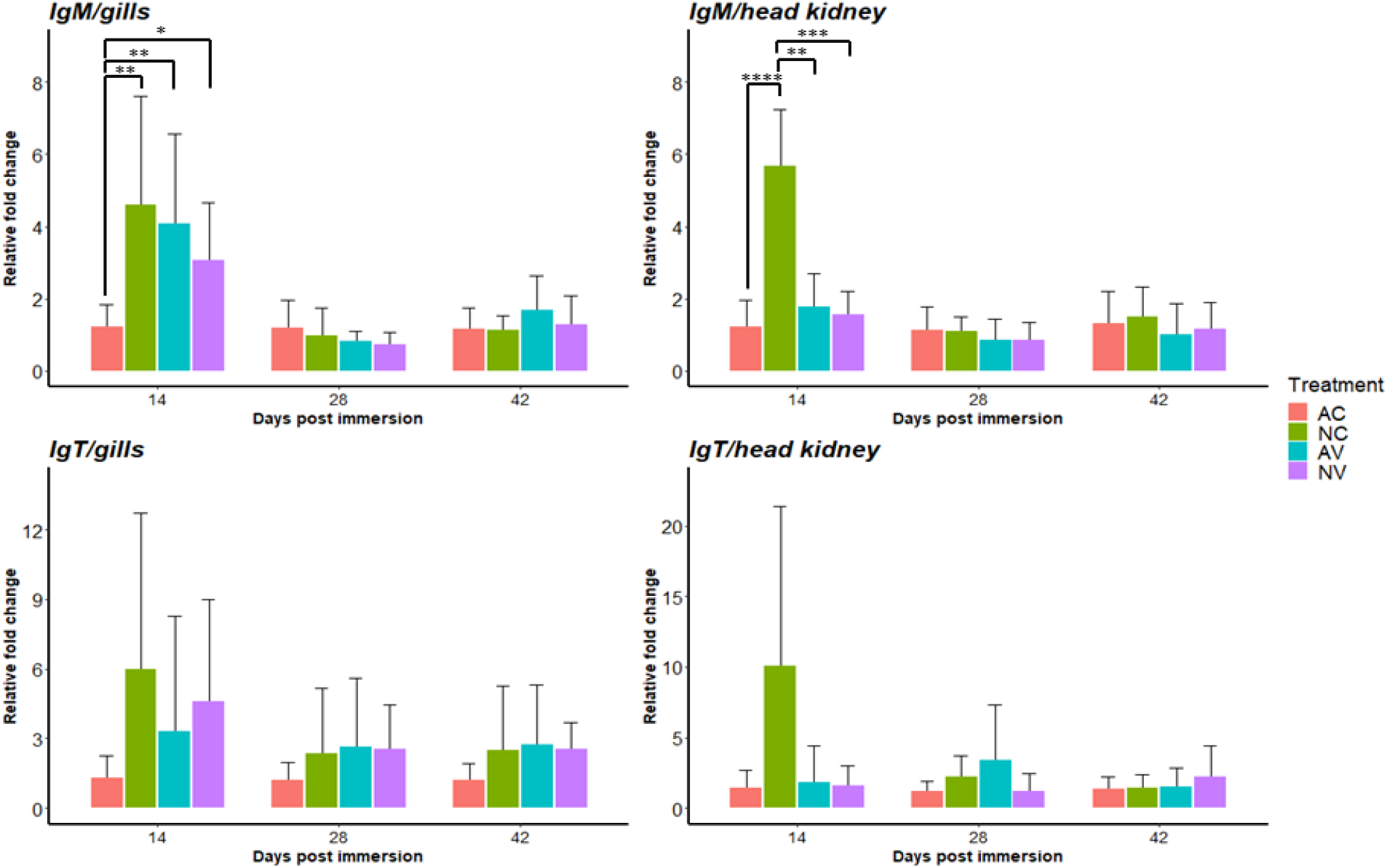
Relative *IgM, IgT* gene expression (Relative fold change ± SD) in the gills and head kidney of Nile tilapia, *Oreochromis niloticus*, post immersion and the first oral booster. Within the same sampling point, the connectors with asterisks indicates a significant difference between two groups using a Dunn’s test (*p* adjust with Bonferroni method). Different levels are (*) *p < 0*.*05*, (**) *p < 0*.*01*, (***) *p < 0*.*001*, (****) *p < 0*.*0001*. Abbreviations: AC = normal aeration control; AV = normal aeration + vaccine; NC = ozone nanobubble treatment control; NV = ozone nanobubble treatment + vaccine.

**Figure 4.**
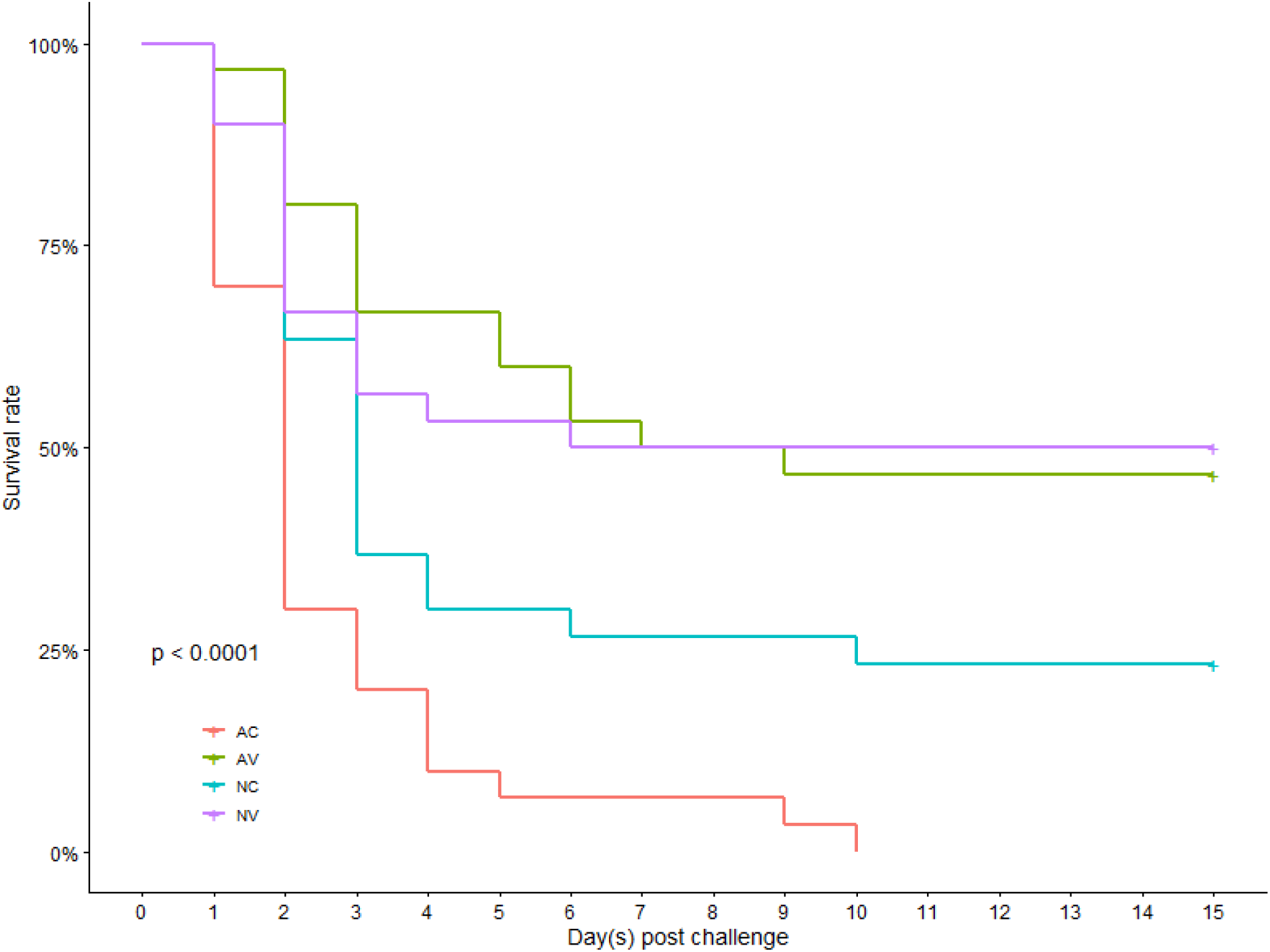
Kaplan-Meier survival rate (%) of Nile tilapia, *Oreochromis niloticus*, post challenge with *Streptococcus agalactiae* (n=30). Abbreviations: AC = normal aeration control; AV = normal aeration + vaccine; NC = ozone nanobubble treatment control; NV = ozone nanobubble treatment + vaccine.

Higher transcript levels in *IgT* expression were recorded in the gills at 14 dpi, however, the expressions were not significantly upregulated in the NC, AV, and NV groups. In the head kidney, the average *IgT* expression was higher only in the NC group, with an approximately ten-fold increase. Nevertheless, no significant differences were observed among the four groups.

### 3.4 Efficacy of the vaccine trial by laboratory scale challenge

Following experimental challenge on day 88 with a high dose of *S. agalactiae* by injection (10^7^ CFU/ fish), fish mortalities began approximately 18 hours post-challenge (Figure 3). Significant mortalities resulted in the first week post-challenge. By day 10 post-challenge, all fish in the AC treatment group had succumbed to infection. At the end of the 15-day challenge period, cumulative mortalities were 100, 76.7, 53.3, and 50.0 % in the AC, NC, AV, and NV groups, respectively. The final corrected relative percent survival (RPS) scores, adjusted by using the AC group, were determined to be 46.7% in the AV group, and 50.0% in the NV group. Moribund fish showed characteristic clinical signs including an impaired, spiraling swimming behavior, skin hemorrhages, and abdominal dropsy. Internally, there was notable enlargement of the liver. Pin head-sized, whitish colonies of *S. agalactiae* were effectively re-isolated on TSA plates from the liver and brain tissues of the challenged fish.

## 4. Discussion

The tilapia industry faces a formidable challenge in the management and control of *S. agalactiae*, which results in substantial economic losses. Given the ongoing crisis of antimicrobial resistance, vaccination has emerged as an essential measure to alleviate the impact of this disease, particularly in the context of tilapia production (Henriksson et al., 2017; Reverter et al., 2020; Schar et al., 2021). Nonetheless, the traditional approach of administering vaccines through injections is regarded as unfavorable by most tilapia producers due to its labor-intensive, time-consuming, potentially costly, and stress-inducing nature. Furthermore, given the comparatively lower market value of tilapia in comparison to other fish species, alternative approaches need to be explored. This study presents an innovative approach for the vaccination of tilapia against *S. agalactiae*, using a combination of three novel strategies, namely an ONb pre-treatment, followed by a “VAC in BAG” - an immersion vaccine given during the transportation phase, and then by “VAC in FEED” - a booster vaccine added to a commercial tilapia diet and presented to fish at two points in the grow-out phase of production. The research builds upon the findings of Linh et al. (2021, 2022) who demonstrated the immuno-activation properties of ONb which enhance antigen uptake during immersion vaccination. Additionally, by recognizing the general reluctance of tilapia producers to vaccinate their fish once they have been stocked into grow-out systems, we propose that the translocation of young fish (fry or fingerlings) from hatcheries to grow-out cages present an opportune time for vaccination. By implementing fish immersion immunization in oxygenated bags (VAC in BAG) during transportation, the effort and infrastructure required for vaccination can be significantly reduced while at the same time minimizing stress because the method negates the need to handle fish. Importantly, our study revealed that a five-hour transportation period, using stocking density of ca. 100 g fish biomass/L, did not result in any mortalities or abnormalities at the point of stocking nor throughout the following seven-days. This indicates that a 5-hour long immersion vaccine during within transport bags is practically safe for the fish of 5 g Nile tilapia.

A vaccine booster is typically given to extend the protection of the fish against a pathogen during production. Once the fish have settled within their culture system, our procedure finds the use of oral boosters (VAC in FEED) more convenient compared to other modes of vaccine delivery (Mutoloki et al., 2015; Yao et al., 2019). Through the integration of both immersion and oral administration methods, the vaccination process can be significantly streamlined. Additionally, the utilization of a heat-killed vaccine further simplifies the vaccine formulation and reduces the reliance on potentially harmful substances like formalin. Our decision to use a heat-killed vaccine in this study aligns with the increasing significance of autogenous vaccines, which can be locally produced using isolates from local outbreaks. This approach enhances accessibility, and offers protection, particularly in situations where commercial vaccines are not readily accessible (Barnes et al., 2022).

Regular monitoring conducted every two weeks revealed the stimulation of the fish’s humoral adaptive immune system by the vaccine, as indicated by specific antibody production. Interestingly, only the NV group exhibited significant levels of anti-*Sa* antibodies during the initial four weeks following immersion. While the AV group required an additional oral booster dose to generate anti-*Sa* antibodies effectively. Consistent with a previous laboratory study conducted by Linh et al. (2022), this field investigation provides further confirmation that pre-treating with ONb can enhance antibody production in tilapia against *S. agalactiae*, potentially through increased antigen uptake and the stimulation of the innate immune response. These findings highlight the promising potential of ONb as a transformative factor in enhancing the immune response of fish during immersion vaccination. Oral boosters were administered during two specific periods, namely 28-32 and 70-72 days post-immersion (dpi) during the grow-out stage. Similar to the research findings from other studies (i.e., Hayat et al., 2021; Yao et al., 2019), the administration of oral boosters has demonstrated the ability to prolong antibody production in fish. Our research shows that there was a correlation with the subsequent rise in antibodies after the first booster; notably those within the NV group which continued to rise until 70 dpi. The AV group, however, only started to show a significant antibody response two weeks after the first oral booster, but not after the primary immersion vaccine. These findings indicate that the oral booster plays a crucial role in eliciting detectable antibody responses in the AV group. In recent laboratory studies, a simple heat-killed *S. agalactiae* vaccine was found to elicit antibody production two weeks after immersion (Linh et al., 2022); a similar response was seen with a nanoparticle vaccine (Ke et al., 2021). As no antibody upregulation was seen in the AV group after the immersion, we assume that this group might go through a priming stage, when memory responses were formed but the adaptive immune system did not express detectable antibodies, after which they then responded strongly to the antigens incorporated into the first VAC in FEED booster. The antibody levels dropped in all vaccinated groups at 84 dpi, despite a second VAC in FEED booster being given at 70 dpi. It is plausible that the shorter three-day period over which the second VAC in FEED booster was given, was not sufficiently long enough to elicit a specific immune response as was seen following the first 5-day VAC in FEED booster.

During our field trial study, it was challenging to eliminate the influence of random factors on the research outcomes. Notably, a considerable number of fish in the unvaccinated groups exhibited positive antibodies against *S. agalactiae*, surpassing the cut-off threshold of 0.05 significance level. This was observed in the non-vaccinated group from 42 to 84 days post-infection (dpi) and in the alternative control group from 56 to 84 dpi. *Streptococcus agalactiae* is a multi-host pathogen commonly found in freshwater (Niu et al., 2020; Delannoy et al., 2021). The experimental cages used in this study were positioned in a ca. 1 hectare pond within which there many fish (i.e., a wild tilapia population having established from escapees from earlier rounds of production), shellfish, and a natural microbial community which may have also included *S. agalactiae*. The field trial was conducted between the months of July to October in Thailand, when water temperatures range typically range from 30 – 32 °C – temperatures which favor the proliferation of *S. agalactiae* (see Kayansamruaj et al., 2014). Following our initial assumption, it appears that the unvaccinated groups developed an antibody response due to their natural exposure to *S. agalactiae* in the pond. Additionally, the unexpected increase in specific antibodies could be attributed to the inadvertent leaching of inactivated bacterial cells during the feed-based oral boosters. The fish in our experiments were grouped and separated by net borders, but water flow might have carried the inactivated bacterial cells to neighboring cages, thereby activating the fish’s immune response through the skin or gill mucosal immune system.

The expression of two important immunoglobulin genes – *IgM* and *IgT*, in two lymphoid organs – the gills and the head kidney, implied that fish’s adaptive immunity was elicited by the immersion vaccine. Immunoglobulin molecules play crucial components in the immune defense system. *IgM* and *IgT* transcripts were detected in the early larval development stages of Nile tilapia (Ke et al., 2021). *IgM* plays a crucial role in adaptive immunity at the systemic level as well as in the mucosal immune response (Salinas et al., 2021). When compared, *IgT* functions are more specialized for mucosal immunity in Nile tilapia (Velázquez et al., 2018).

In both vaccinated fish groups, the *IgM* expression was significantly higher in the gills at 14 dpi, while it was upregulated in the head kidney of only a small number of the fish that were sampled indicting it is not significantly upregulated. A similar expression pattern was observed in the *IgT* gene of both organs. These observations could potentially reflect the varied responses of individuals when administered the immersion vaccine. Unexpectedly, the group that received the ONb treatment without vaccination exhibited the highest levels of *IgM* and *IgT* expression in both lymphoid organs. Previous research has demonstrated that ONb can modulate both the innate and adaptive immune systems in tilapia (Dien et al., 2021; Linh et al., 2021, 2022). Our fish were raised in an open environment that harbored a diverse array of microorganisms, including other potential pathogens of Nile tilapia. It is likely that the immune-modulating effects of ONb stimulated the fish’s immunity towards other unknown microorganisms present in the water upon stocking. Additionally, the expression of *IgM* and *IgT* in the head kidney of the non-vaccinated (NC) fish was significantly higher when compared to both the vaccinated groups (AV and NV). We speculate that the vaccinated groups exhibited early and pronounced upregulation of *IgM* and *IgT* transcripts following vaccination, which may have occurred before the first 14-day post-infection (dpi) evaluation point. In contrast, the expression in the non-vaccinated (NC) group was delayed until their exposure to microbes in the water. Consequently, the peak expression in this group was observed at the 14-day post-infection (dpi) time point.

Following the administration of the first VAC in FEED booster, no significant upregulation was observed in any of the groups. It is possible that the immune response occurred in other lymphoid organs, i.e., in others, other than the gills and head kidney where transcripts were determined. Nevertheless, we acknowledge that our understanding of the complete immune response post-vaccination is limited due to the wide spatial distribution of sampling points and the absence of investigations into the gut-associated lymphoid organ following oral boosters.

Even though a challenge test was conducted approximately three months after vaccination, after which the antibody level had dwindled, our vaccination program exhibited a moderate level of protection at 50% in the NV group that fish was treated with ONb. The protection conferred by the vaccination strategy in the NV group was not notably higher compared to the normal vaccinated AV group at 46.7%. In many vaccine studies for tilapia, the congruence between antibody response and vaccine efficacy by was evident (Kitiyodom et al., 2019; Linh et al., 2022; Mai et al., 2021). As antibody had dropped to statistically similar levels in both vaccinated groups four days before the challenge conduction, it may explain the negligible difference between those groups in the current study. Nevertheless, as the fish in the NV group exhibited a more robust antibody response within the initial 70 days of the study, they may display greater resilience when exposed to the pathogen earlier, indicating potential endurance throughout the course of infection.

Several field trial vaccines have been utilized to safeguard tilapia against *S. agalactiae*, with reported vaccine efficacies based on endemic survival rates (i.e., fish survival is based on monitoring in the field rather than from lab-based experimental challenges). For instance, an injectable vaccine targeting serotype Ia and III demonstrated protection levels exceeding 77% when administered in double doses during the grow-out phase from 25-30 g to 900-1000 g fish (Kannika et al., 2022). Similarly, a double-dose, feed-based vaccine was found to result in a RPS of 75% after a 16-week trial (Ismail et al., 2017). In contrast to the aforementioned examples, our trial included a challenge test where live *S. agalactiae* was injected into a small portion of the population. In fish vaccine studies, injection ensures that all tested fish receive consistent amounts of the pathogen. Injecting live pathogens, however, can be more detrimental to fish since it bypasses the protective epithelial barrier, which plays a crucial role in preventing infections in the first place. As a result, an injection challenge would likely lead to lower survival rates, as observed in other field vaccine trials. Nonetheless, the decision to utilize an experimental injection challenge ensured that mortalities were primarily caused by *S. agalactiae*, as co-infections in natural conditions often contribute significantly to mortality rates in tilapia aquaculture (Dong et al., 2015).

Interestingly, the NC group had a higher survival rate than the AC group, despite receiving no vaccine. In lab-scale experiments previously, the ONb pre-treatment could enhance the immunity of the tilapia subsequently enabling them to have a better survival under infection (Dien et al., 2021; Linh et al., 2021). We hypothesize that ONb application in the NC group had excited the fish’s immune system before stocking so that they were better able to mount specific immunity via natural exposure to live *S. agalactiae* in the pond environment. The unexpected significant upregulation of this group, when compared with the control AC group, as mentioned above may also support this hypothesis. Furthermore, it is possible that the immunomodulatory effect of ONb played a role in stimulating the innate immune response of the fish. However, it remains an area of interest for further investigation to determine the duration of this effect and whether it can provide protection to the fish when they encounter an infection.

In summary, this study presents an innovative approach to vaccinating Nile tilapia against *S. agalactiae* infection. The approach integrates ONb pre-treatment to activate the fish immune system, followed by a novel immersion vaccination method known as VAC in BAG during the transportation of fish from the hatchery to the grow-out farm. The fish are then supplemented with oral boosters (VAC in FEED). This novel vaccination strategy has demonstrated effectiveness in enhancing the specific immune response and in extending the duration of anti-*Sa* antibodies for approximately three months. Moreover, vaccinated fish near the end of the culture cycle exhibited a satisfactory level of protection when challenged with a high dose of *S. agalactiae*. Collectively, this innovative vaccination strategy holds promise as a transformative alternative to the laborious and expensive injection-based mode of vaccination that is currently used. This novel and innovative vaccination regimen empowers Nile tilapia producers with cost-effective alternatives for protecting their stocks, addressing a current lack of effective protective measures in place.

## Acknowledgements

This research project received financial support from the UK government - Department of Health and Social Care (DHSC), Global AMR Innovation Fund (GAMRIF), and the International Development Research Center (IDRC), Ottawa, Canada.

## Data availability

The available data has been presented in this manuscript.

## Author contributions

**Nguyen Tien Vinh**: Investigation, Methodology, Formal analysis, Writing – original draft, Software, and Resources. **Ha Thanh Dong**: Conceptualization, Data curation, Writing – review & editing, Supervision, Validation, Funding acquisition, and Project administration. **Saengchan Senapin**: Conceptualization, Data curation, Writing – review & editing. **Suntree Pumpuang**: Investigation. **Nguyen Giang Thu Lan**: Investigation, Methodology, and Writing – review & editing. **Bulakorn Wilairat**: Investigation and Methodology. **Pradnya R. Garud**: Investigation and Methodology. **Sophie St-Hilaire**: Writing – review & editing, and Funding acquisition. **Nguyen Vu Linh**: Methodology, and Writing – review & editing. **Wattana Phanphut:** Writing – review & editing. **Andrew P. Shinn**: Writing – review & editing.

## Supplemental data

**Figure S1.**
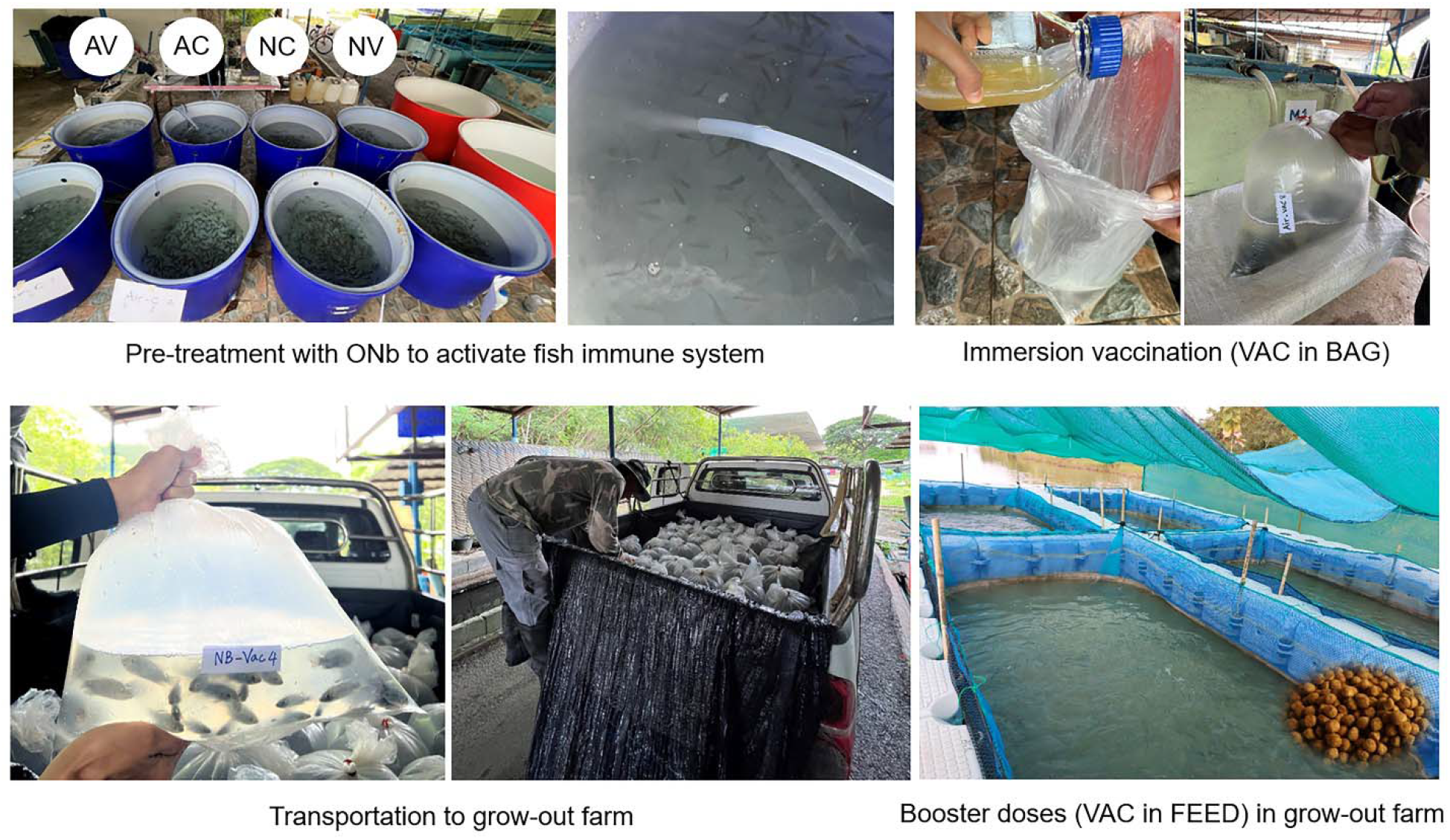
Actual activities during the ozone nanobubble (ONb) pre-treatment, VAC in BAG and VAC in FEED field trial. Abbreviations: AC = normal aeration control; AV = normal aeration + vaccine; NC = ozone nanobubble treatment control; NV = ozone nanobubble treatment + vaccine.

## References

Abu-Elala, N. M., Samir, A., Wasfy, M., & Elsayed, M. (2019). Efficacy of Injectable and Immersion Polyvalent Vaccine against Streptococcal Infections in Broodstock and Offspring of Nile tilapia (Oreochromis niloticus). Fish & Shellfish Immunology, 88, 293–300. https://doi.org/10.1016/J.FSI.2019.02.042

Adams, A. (2019). Progress, challenges and opportunities in fish vaccine development. Fish & Shellfish Immunology, 90, 210–214. https://doi.org/10.1016/J.FSI.2019.04.066

Amal, M. N. A., Saad, M. Z., Zahrah, A. S., & Zulkafli, A. R. (2015). Water quality influences the presence of Streptococcus agalactiae in cage cultured red hybrid tilapia, Oreochromis niloticus × Oreochromis mossambicus. Aquaculture Research, 46(2), 313–323. https://doi.org/10.1111/ARE.12180

Barnes, A. C., Silayeva, O., Landos, M., Dong, H. T., Lusiastuti, A., Phuoc, L. H., & Delamare-Deboutteville, J. (2022). Autogenous vaccination in aquaculture: A locally enabled solution towards reduction of the global antimicrobial resistance problem. Reviews in Aquaculture, 14(2), 907–918. https://doi.org/10.1111/RAQ.12633

Bøgwald, J., & Dalmo, R. A. (2019). Review on Immersion Vaccines for Fish: An Update 2019. Microorganisms 2019, Vol. 7, Page 627, 7(12), 627. https://doi.org/10.3390/MICROORGANISMS7120627

Brudeseth, B. E., Wiulsrød, R., Fredriksen, B. N., Lindmo, K., Løkling, K. E., Bordevik, M., Steine, N., Klevan, A., & Gravningen, K. (2013). Status and future perspectives of vaccines for industrialised fin-fish farming. Fish & Shellfish Immunology, 35(6), 1759–1768. https://doi.org/10.1016/J.FSI.2013.05.029

Chettri, J. K., Jaafar, R. M., Skov, J., Kania, P. W., Dalsgaard, I., & Buchmann, K. (2015). Booster immersion vaccination using diluted Yersinia ruckeri bacterin confers protection against ERM in rainbow trout. Aquaculture, 440, 1–5. https://doi.org/10.1016/J.AQUACULTURE.2015.01.027

Chitmanat, C., Lebel, P., Whangchai, N., Promya, J., & Lebel, L. (2016). Tilapia diseases and management in river-based cage aquaculture in northern Thailand. Journal of Applied Aquaculture, 28(1), 9–16. https://doi.org/10.1080/10454438.2015.1104950

Delannoy, C. M. J., Samai, H., & Labrie, L. (2021). Streptococcus agalactiae serotype IV in farmed tilapia. Aquaculture, 544, 737033. https://doi.org/10.1016/J.AQUACULTURE.2021.737033

Dien, L. T., Linh, N. V., Sangpo, P., Senapin, S., St-Hilaire, S., Rodkhum, C., & Dong, H. T. (2021). Ozone nanobubble treatments improve survivability of Nile tilapia (Oreochromis niloticus) challenged with a pathogenic multi-drug-resistant Aeromonas hydrophila. Journal of Fish Diseases, 44(9), 1435–1447. https://doi.org/10.1111/JFD.13451

Dong, H. T., Nguyen, V. V., Le, H. D., Sangsuriya, P., Jitrakorn, S., Saksmerprome, V., Senapin, S., & Rodkhum, C. (2015). Naturally concurrent infections of bacterial and viral pathogens in disease outbreaks in cultured Nile tilapia (Oreochromis niloticus) farms. Aquaculture, 448, 427–435. https://doi.org/10.1016/J.AQUACULTURE.2015.06.027

FAO. (2020). The State of World Fisheries and Aquaculture 2020. Sustainability in action. https://doi.org/10.4060/ca9229en

Frey, A., di Canzio, J., & Zurakowski, D. (1998). A statistically defined endpoint titer determination method for immunoassays. Journal of Immunological Methods, 221(1–2), 35–41. https://doi.org/10.1016/S0022-1759(98)00170-7

Georgiadis, M. P., Gardner, I. A., & Hedrick, R. P. (2001). The role of epidemiology in the prevention, diagnosis, and control of infectious diseases of fish. Preventive Veterinary Medicine, 48(4), 287–302. https://doi.org/10.1016/S0167-5877(00)00202-6

Hayat, M., Yusoff, M. S. M., Samad, M. J., Razak, I. S. A., Yasin, I. S. M., Thompson, K. D., & Hasni, K. (2021). Efficacy of Feed-Based Formalin-Killed Vaccine of Streptococcus iniae Stimulates the Gut-Associated Lymphoid Tissues and Immune Response of Red Hybrid Tilapia. Vaccines, 9(1), 51. https://doi.org/10.3390/VACCINES9010051

Henriksson, P. J. G., Rico, A., Troell, M., Klinger, D. H., Buschmann, A. H., Saksida, S., Chadag, M. V., & Zhang, W. (2017). Unpacking factors influencing antimicrobial use in global aquaculture and their implication for management: a review from a systems perspective. Sustainability Science, 13(4), 1105–1120. https://doi.org/10.1007/S11625-017-0511-8

Ismail, M. S., Syafiq, M. R., Siti-Zahrah, A., Fahmi, S., Shahidan, H., Hanan, Y., Amal, M. N. A., & Zamri Saad, M. (2017). The effect of feed-based vaccination on tilapia farm endemic for streptococcosis. Fish & Shellfish Immunology, 60, 21–24. https://doi.org/10.1016/J.FSI.2016.11.040

Jansen, K. U., & Anderson, A. S. (2018). The role of vaccines in fighting antimicrobial resistance (AMR). Human Vaccines & Immunotherapeutics, 14(9), 2142–2149. https://doi.org/10.1080/21645515.2018.1476814

Jantrakajorn, S., Maisak, H., & Wongtavatchai, J. (2014). Comprehensive Investigation of Streptococcosis Outbreaks in Cultured Nile Tilapia, Oreochromis niloticus, and Red Tilapia, Oreochromis sp., of Thailand. Journal of the World Aquaculture Society, 45(4), 392–402. https://doi.org/10.1111/JWAS.12131

Jhunkeaw, C., Khongcharoen, N., Rungrueng, N., Sangpo, P., Panphut, W., Thapinta, A., Senapin, S., St-Hilaire, S., & Dong, H. T. (2021). Ozone nanobubble treatment in freshwater effectively reduced pathogenic fish bacteria and is safe for Nile tilapia (Oreochromis niloticus). Aquaculture, 534, 736286. https://doi.org/10.1016/J.AQUACULTURE.2020.736286

Kannika, K., Sirisuay, S., Kondo, H., Hirono, I., Areechon, N., & Unajak, S. (2022). Trial Evaluation of Protection and Immunogenicity of Piscine Bivalent Streptococcal Vaccine: From the Lab to the Farms. Vaccines, 10(10), 1625. https://doi.org/10.3390/VACCINES10101625

Kassambara, A. (2023). rstatix: Pipe-Friendly Framework for Basic Statistical Tests (R package version 0.7.2). https://rpkgs.datanovia.com/rstatix/authors.html

Kassambara, A., Kosinski, M., Biecek, P., & Fabian, S. (2021). Package “survminer” (0.4.9). https://cran.r-project.org/web/packages/survminer/index.html

Kayansamruaj, P., Pirarat, N., Hirono, I., & Rodkhum, C. (2014). Increasing of temperature induces pathogenicity of Streptococcus agalactiae and the up-regulation of inflammatory related genes in infected Nile tilapia (Oreochromis niloticus). Veterinary Microbiology, 172(1–2), 265–271. https://doi.org/10.1016/J.VETMIC.2014.04.013

Ke, X., Liu, Z., Chen, S., Chen, Z., Zhang, D., Gao, F., & Lu, M. (2021). The immune efficacy of a Streptococcus agalactiae immersion vaccine for different sizes of young tilapia. Aquaculture, 534, 736289. https://doi.org/10.1016/J.AQUACULTURE.2020.736289

Kitiyodom, S., Kaewmalun, S., Nittayasut, N., Suktham, K., Surassmo, S., Namdee, K., Rodkhum, C., Pirarat, N., & Yata, T. (2019). The potential of mucoadhesive polymer in enhancing efficacy of direct immersion vaccination against Flavobacterium columnare infection in tilapia. Fish & Shellfish Immunology, 86, 635–640. https://doi.org/10.1016/J.FSI.2018.12.005

Linh, N. V., Dien, L. T., Panphut, W., Thapinta, A., Senapin, S., St-Hilaire, S., Rodkhum, C., & Dong, H. T. (2021). Ozone nanobubble modulates the innate defense system of Nile tilapia (Oreochromis niloticus) against Streptococcus agalactiae. Fish and Shellfish Immunology, 112, 64–73. https://doi.org/10.1016/j.fsi.2021.02.015

Linh, N. V., Dien, L. T., Sangpo, P., Senapin, S., Thapinta, A., Panphut, W., St-Hilaire, S., Rodkhum, C., & Dong, H. T. (2022). Pre-treatment of Nile tilapia (Oreochromis niloticus) with ozone nanobubbles improve efficacy of heat-killed Streptococcus agalactiae immersion vaccine. Fish and Shellfish Immunology, 123, 229–237. https://doi.org/10.1016/j.fsi.2022.03.007

Livak, K. J., & Schmittgen, T. D. (2001). Analysis of relative gene expression data using real-time quantitative PCR and the 2(-Delta Delta C(T)) Method. Methods (San Diego, Calif.), 25(4), 402–408. https://doi.org/10.1006/METH.2001.1262

Ma, J., Bruce, T. J., Jones, E. M., & Cain, K. D. (2019). A Review of Fish Vaccine Development Strategies: Conventional Methods and Modern Biotechnological Approaches. Microorganisms, 7(11), 569. https://doi.org/10.3390/MICROORGANISMS7110569

Mai, T. T., Kayansamruaj, P., Taengphu, S., Senapin, S., Costa, J. Z., del-Pozo, J., Thompson, K. D., Rodkhum, C., & Dong, H. T. (2021). Efficacy of heat-killed and formalin-killed vaccines against Tilapia tilapinevirus in juvenile Nile tilapia (Oreochromis niloticus). Journal of Fish Diseases, 44(12), 2097–2109. https://doi.org/10.1111/JFD.13523

Mian, G. F., Godoy, D. T., Leal, C. A. G., Yuhara, T. Y., Costa, G. M., & Figueiredo, H. C. P. (2009). Aspects of the natural history and virulence of S. agalactiae infection in Nile tilapia. Veterinary Microbiology, 136(1–2), 180–183. https://doi.org/10.1016/J.VETMIC.2008.10.016

Miccoli, A., Manni, M., Picchietti, S., & Scapigliati, G. (2021). State-of-the-Art Vaccine Research for Aquaculture Use: The Case of Three Economically Relevant Fish Species. Vaccines, 9(2), 140. https://doi.org/10.3390/VACCINES9020140

Micoli, F., Bagnoli, F., Rappuoli, R., & Serruto, D. (2021). The role of vaccines in combatting antimicrobial resistance. Nature Reviews Microbiology 2021 19:5, 19(5), 287–302. https://doi.org/10.1038/s41579-020-00506-3

Mondal, H., & Thomas, J. (2022). A review on the recent advances and application of vaccines against fish pathogens in aquaculture. Aquaculture International 2022 30:4, 30(4), 1971–2000. https://doi.org/10.1007/S10499-022-00884-W

Mutoloki, S., Munang’andu, H. M., & Evensen, &#x216;. (2015). Oral vaccination of fish - antigen preparations, uptake, and immune induction. Frontiers in Immunology, 6(OCT), 519. https://doi.org/10.3389/FIMMU.2015.00519/BIBTEX

Niu, G., Khattiya, R., Zhang, T., Boonyayatra, S., & Wongsathein, D. (2020). Phenotypic and genotypic characterization of Streptococcus spp. isolated from tilapia (Oreochromis spp.) cultured in river-based cage and earthen ponds in Northern Thailand. Journal of Fish Diseases, 43(3), 391–398. https://doi.org/10.1111/JFD.13137

Plumb, J. A., & Vinitnantharat, S. (1993). Vaccination of channel catfish, Ictalurus punctatus (Rafinesque), by immersion and oral booster against Edwardsiella ictaluri. Journal of Fish Diseases, 16(1), 65–71. https://doi.org/10.1111/J.1365-2761.1993.TB00848.X

Reverter, M., Sarter, S., Caruso, D., Avarre, J. C., Combe, M., Pepey, E., Pouyaud, L., Vega-Heredía, S., de Verdal, H., & Gozlan, R. E. (2020). Aquaculture at the crossroads of global warming and antimicrobial resistance. Nature Communications 2020 11:1, 11(1), 1–8. https://doi.org/10.1038/s41467-020-15735-6

Romana-Eguia, M. R. R., Eguia, R. v., & Pakingking, Jr. R. V. (2020). Tilapia culture: The basics. Aquaculture Department, Southeast Asian Fisheries Development Center. http://hdl.handle.net/10862/5842

Salinas, I. Fernández-Montero Á., Ding, Y., & Sunyer, J. O. (2021). Mucosal immunoglobulins of teleost fish: A decade of advances. Developmental and Comparative Immunology, 121, 104079. https://doi.org/10.1016/j.dci.2021.104079

Schar, D., Zhao, C., Wang, Y., Larsson, D. G. J., Gilbert, M., & Van Boeckel, T. P. (2021). Twenty-year trends in antimicrobial resistance from aquaculture and fisheries in Asia. Nature Communications 2021 12:1, 12(1), 1–10. https://doi.org/10.1038/s41467-021-25655-8

Schmidt, J. G., Henriksen, N. H., & Buchmann, K. (2016). ERM booster vaccination of rainbow trout using diluted bacterin: Field studies. Aquaculture, 464, 262–267. https://doi.org/10.1016/J.AQUACULTURE.2016.07.001

Shinn, A., Dong, H., Vinh, N., Wongwaradechkul, R., & Lio-Po, G. (2023). Infectious Diseases of Warmwater Fish in Fresh Water. In Climate Change on Diseases and Disorders of Finfish in Cage Culture (pp.202–277). https://doi.org/10.1079/9781800621640.0006

Therneau, T. (2023). A Package for Survival Analysis in R (R package version 3.5-0). https://CRAN.R-project.org/package=survival

Velázquez, J., Acosta, J., Lugo, J. M., Reyes, E., Herrera, F., González, O., Morales, A., Carpio, Y., & Estrada, M. P. (2018). Discovery of immunoglobulin T in Nile tilapia (Oreochromis niloticus): A potential molecular marker to understand mucosal immunity in this species. Developmental & Comparative Immunology, 88, 124–136. https://doi.org/10.1016/J.DCI.2018.07.013

Wickham, H. (2016). ggplot2: Elegant Graphics for Data Analysis. Springer-Verlag New York. https://ggplot2.tidyverse.org

Wickham, H., François, R., Henry, L., & Müller, K. (2022). dplyr: A Grammar of Data Manipulation (1.0.10). https://dplyr.tidyverse.org, https://github.com/tidyverse/dplyr.

Yao, Y. Y., Chen, D. D., Cui, Z. W., Zhang, X. Y., Zhou, Y. Y., Guo, X., Li, A. H., & Zhang, Y. A. (2019). Oral vaccination of tilapia against Streptococcus agalactiae using Bacillus subtilis spores expressing Sip. Fish & Shellfish Immunology, 86, 999–1008. https://doi.org/10.1016/J.FSI.2018.12.060

Zhu, L., Yang, Q., Huang, L., Wang, K., Wang, X., Chen, D., Geng, Y., Huang, X., Ouyang, P., & Lai, W. (2017). Effectivity of oral recombinant DNA vaccine against Streptococcus agalactiae in Nile tilapia. Developmental & Comparative Immunology, 77, 77–87. https://doi.org/10.1016/J.DCI.2017.07.024

